# Realistic *in silico* generation and augmentation of single cell RNA-seq data using Generative Adversarial Neural Networks

**DOI:** 10.1101/390153

**Authors:** Mohamed Marouf, Pierre Machart, Vikas Bansal, Christoph Kilian, Daniel S. Magruder, Christian F. Krebs, Stefan Bonn

**Affiliations:** Institute of Medical Systems Biology, University Medical Center Hamburg-Eppendorf, Germany; German Center for Neurodegenerative Diseases Tuebingen, Germany; Genevention GmbH, Goettingen, Germany; Center for Internal Medicine, III. Medical Clinic and Polyclinic, University Medical Center Hamburg-Eppendorf, Germany

**Keywords:** Deep Learning, Generative Adversarial Neural Networks, Machine Learning, data augmentation, single cell RNA-seq, RNA sequencing, simulation

## Abstract

A fundamental problem in biomedical research is the low number of observations available, mostly due to a lack of available biosamples, prohibitive costs, or ethical reasons. Augmenting few real observations with generated *in silico* samples could lead to more robust analysis results and a higher reproducibility rate. Here we propose the use of conditional single cell Generative Adversarial Neural Networks (cscGANs) for the realistic generation of single cell RNA-seq data. cscGANs learn non-linear gene-gene dependencies from complex, multi cell type samples and use this information to generate realistic cells of defined types. Augmenting sparse cell populations with cscGAN generated cells improves downstream analyses such as the detection of marker genes, the robustness and reliability of classifiers, the assessment of novel analysis algorithms, and might reduce the number of animal experiments and costs in consequence. cscGANs outperform existing methods for single cell RNA-seq data generation in quality and hold great promise for the realistic generation and augmentation of other biomedical data types.

## Introduction

Biological systems are usually highly complex, as intra-and inter-cellular communication, for example, are orchestrated via the non-linear interplay of tens to hundreds of thousands of different molecules^1^. Recent technical advances have enabled scientists to scrutinize these complex interactions, measuring the expression of thousands of genes at the same time, for instance^2^. Unfortunately, this complexity often becomes a major hurdle as the number of observations can be relatively small, due to economical or ethical considerations or simply because the number of available patient samples is low. Next to technically-induced measurement biases, this problem of too few observations in the face of many parameters might be one of the most prominent bottlenecks in biomedical research^1^. Thus, limited numbers of observations can lead to sampling bias that could reduce the reproducibility of experimental results, a well-known problem in biomedicine.

While the number of biological samples might be limited, realistic *in silico* generation of observations could accommodate for this unfavorable setting. In practice, *in silico* generation has seen success in computer vision when used for ‘data augmentation’, whereby *in silico* generated samples are used alongside the original ones to artificially increase the number of observations^3^. While classically realistic data modeling relies on a thorough understanding of the laws underlying the production of such data, current methods of choice for photo-realistic image generation rely on Deep Learning-based Generative Adversarial Networks (GANs)^4–7^ and Variational Autoencoders (VAEs)^8,9^.

GANs involve a generator that learns to output realistic *in silico* generated samples, and a critic that learns to spot the differences between real samples and generated ones (Fig. S1). An ‘adversarial’ training procedure allows for those two Neural Networks to compete against each other in a mutually beneficial way.

While data augmentation has been a recent success story in various fields of computer science, the development and usage of GANs and VAEs for omics data augmentation has yet to be achieved. As a proof of concept that realistic *in silico* generation can be applied to biomedical omics data, we focus on the generation of single cell RNA (scRNA) sequencing data. scRNA sequencing has made it possible to evaluate genome-wide gene expression of thousands to millions of cells in a single experiment^10^. This detailed information across genes and cells opens the door to a much deeper understanding of cell type heterogeneity in a tissue, cell differentiation, and cell type-specific disease etiology.

In this manuscript, we establish how a single cell Generative Adversarial Network (scGAN) can be leveraged to generate realistic scRNA-seq data. We further demonstrate that our scGAN can use conditioning (cscGAN) to produce specific cell types or sub-populations, on-demand. Finally, we show how our models can successfully augment sparse datasets to improve the quality and significance of downstream statistical analyses. To the best of our knowledge, this constitutes the first attempt to apply these groundbreaking methods for the augmentation of sequencing data.

## Results

### Realistic generation of scRNA-seq data using an scGAN

Given the great success of GANs in producing photorealistic images, we hypothesize that similar approaches could be used to generate realistic scRNA-seq data. Generated scRNA-seq data that augments experimental data could improve downstream analyses such as classification and the testing of novel algorithms, as briefly outlined in the introduction. To distinguish experimental scRNA-seq data from data produced by GANs we will use the terms ‘real’ and ‘generated’ cells, respectively.

To build and evaluate different GAN models for scRNA-seq data generation we used a Peripheral Blood Mononuclear Cell (PBMC) scRNA-seq dataset with 68,579 cells^11^ (Table S1, Methods). The PBMC dataset contains many distinct immune cell types, which yield clear clusters that can be assigned their cell type identity with marker genes (genes specifically expressed in a cluster). The aforementioned features of the PBMC dataset make it ideal for the evaluation of our scGAN performance.

Since it is notoriously difficult to evaluate the quality of generative models^12^, we used three evaluation criteria inspired by single cell data analysis: t-SNE, marker gene correlation, and classification performance (see Methods for evaluation details). These metrics are used as quantitative and qualitative measures to assess the synthesized cells. Based on these criteria, the best performing single cell GAN (scGAN) model was a GAN minimizing the Wasserstein distance^13^, relying on two Fully-Connected Neural Networks with batch normalization (Fig. S1). We found that the quality of the generated cells greatly improved when the training cells were scaled to exhibit a constant total count of 20.000 reads per cell. In addition to this pre-processing step, we added a custom Library Size Normalization (LSN) function to our scGAN’s generator so that it explicitly outputs generated cells with a total read count equal to that of the training data (20,000 reads per cell) (Fig. S1). Our LSN function greatly improved training speed and stability and gave rise to the best performing models based on the aforementioned metrics. Further details of the model selection and (hyper)-parameter optimization can be found in the methods section.

For a qualitative assessment of the results, we used t-SNE^14,15^ to obtain a two-dimensional representation of generated and real cells from the test set (Fig. 1A, Fig. S2). The scGAN generates cells that represent every cluster of the data it was trained on and the expression patterns of marker genes are accurately learnt by scGAN (Fig. S3).

**Figure 1.**
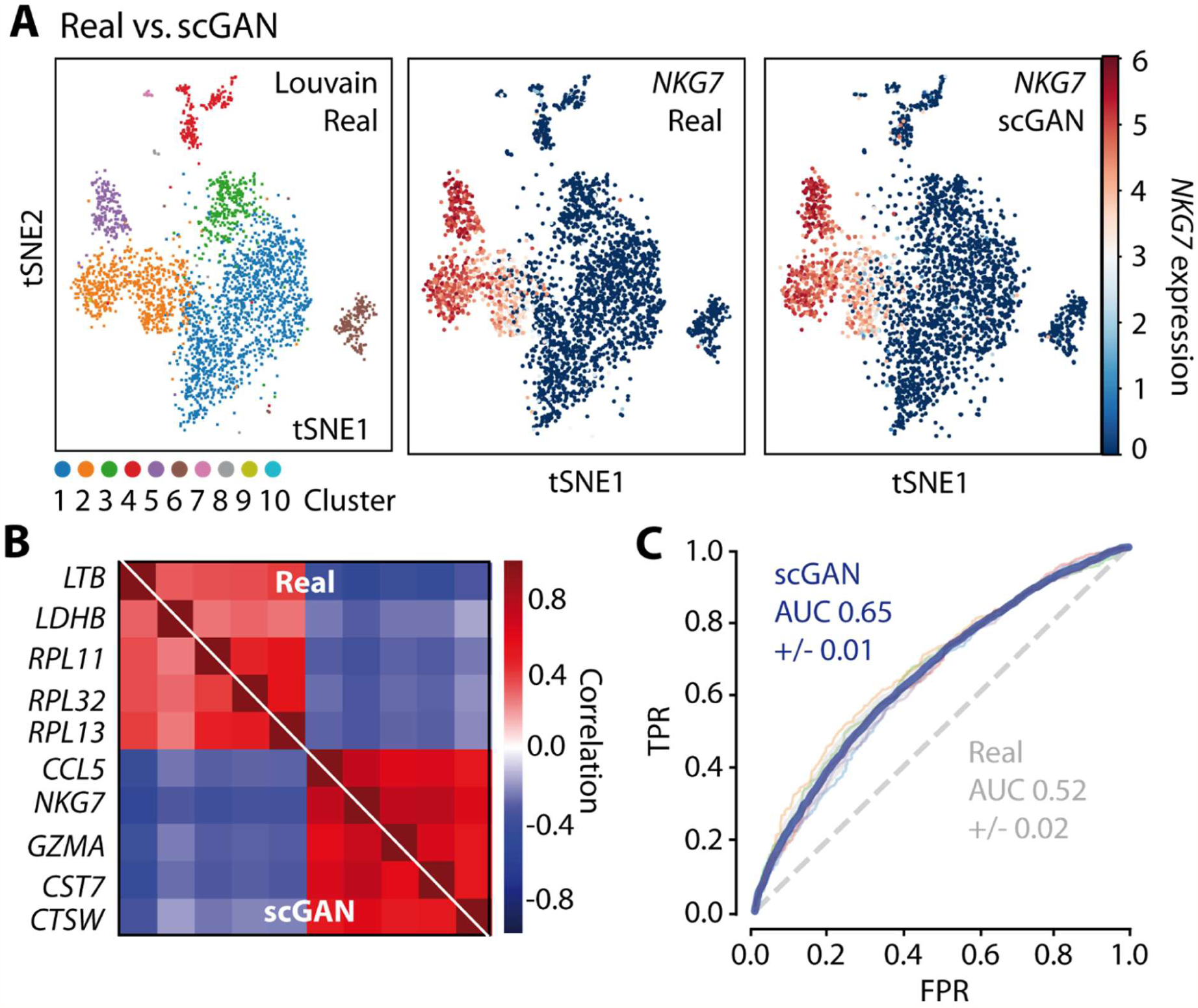
Evaluation of the scGAN generated PBMC cells. A: t-SNE visualization of the Louvain-clustered real cells (left) and the NKG7 gene expression in real (middle) and scGAN generated (right) cells. B: Pearson correlation of marker genes for the scGAN generated (bottom left) and the real (upper right) data. C: Cross validation ROC curve of a RF classifying real and simulated cells (scGAN in blue, chance-level in gray).

Furthermore, the scGAN is able to model inter gene dependencies and correlations, which are a hallmark of biological gene-regulatory networks. To prove this point we compared the correlation and distribution of the counts of cluster-specific marker genes (Fig. 1B) and 100 highly variable genes between generated and real cells (Fig. S4). While the scGAN captures the gene count distributions and correlations well, other state-of-the-art scRNA-seq simulation tools like Splatter^16^ do not model cluster-specific gene correlations appropriately (Fig. S4). While Splatter models some marginal distribution of the read counts (Fig. S5), they struggle to learn the joint distribution of these counts (cluster-specific gene dependencies) (Fig. S4). This is further corroborated by the fact that t-SNE visualizations of Splatter generated data show one homogenous population of cells instead of the different subpopulations of the real and scGAN generated data (Fig. S4). The co-regulation of genes is, however, an important aspect of biological gene-regulatory networks and should be taken into account when generating *in silico* data to test novel analysis methods^17^.

To show that the scGAN generates realistic cells, we trained a Random Forest (RF) classifier^18^ to distinguish between real and generated data. The hypothesis is that a classifier should have a (close to) chance-level performance when the generated and real data are highly similar. Indeed the RF classifier only reaches 0.65 Area Under the Curve (AUC) when discriminating between the real cells and the scGAN generated data (blue curve in Fig. 1C) and 0.52 AUC when tasked to distinguish real from real data (positive control).

The results from the classification, marker gene correlation, and t-SNE corroborate that the scGAN generates realistic data from complex distributions, outperforming existing methods for *in silico* scRNA-seq data generation.

### scGAN runtime scales sub-linearly with the number of cells

Motivated by the ever-increasing number of cells in scRNA-seq datasets, we next wanted to assess how the scGAN training time scales with the number of cells and data complexity (e.g. number of genes and cell types). We therefore trained the scGAN on the currently largest published scRNA-seq data set consisting of 1.3 million mouse brain cells and measured both the time and performance of the model with respect to the number of cells used (Table S1, Fig. S6).

Interestingly, while the best scGAN for the 68k PBMC dataset was trained for over 10,000 epochs, satisfactory results for the 1.3M mouse brain dataset were obtained when training for “only” 1,500 epochs. This suggests that the runtime to accurately train an scGAN could be scaling sub-linearly with respect to the number of training cells. In this context it should be noted that the actual time required to train an scGAN depends on the data size and complexity and on the computer architecture used, necessitating at least one high-performance GPU card.

Our results indicate that the scGAN performs consistently well on different scRNA-seq datasets with varying complexity and size, learning realistic representations of millions of cells.

### Conditional generation of specific cell types

scRNA-seq *in silico* data generation reaches its full potential when specific cells of interest could be generated on demand, which is not possible with the scGAN model. This conditional generation of cell types could be used to increase the number of a sparse, specific population of cells that might represent only a small fraction of the total cells sequenced.

To generate specific cell types of interest while learning multi-cell type complex data, we developed and evaluated various conditional scGAN architectures. Common to all these models is that the conditional scGAN learns to generate cells of specific types while being trained on the complete multi cell type dataset. The cell type information is then associated to the genes’ expression values of each cell during the training. These tags can then be used to generate scRNA-seq data of a specific type, in our case of specific cluster of PBMCs. The best performing conditional scGAN model (cscGAN) utilized a projection discriminator^19^, along with Conditional Batch Normalization^20^ and an LSN function in the generator. Again, model selection and optimization details can be found in the methods.

Model performance was assessed on the PBMC dataset using t-SNE, marker gene correlation, and classification. The cscGAN learns the complete distribution of clusters of the PBMC data and can conditionally represent each of the ten Louvain clusters on demand. The t-SNE results for the conditional generation of cluster 2 and cluster 6 cells are shown in Fig. 2A and Fig. 2B, respectively. Figure 2A highlights the real (red, left and middle) and generated (blue, left and right) cells for cluster 2, while the real cells of all other clusters are shown in grey. The cscGAN generates cells that are overlapping with the real cluster of interest in the t-SNE visualizations. In addition, the cscGAN also accurately captures inter-and intra-cluster gene-gene dependencies as visualized in the marker gene correlation plots in Fig. S 7. The assumption that the cscGAN generates conditional cells that are very similar to the real cells of the cluster of interest is substantiated in the final classification task. A RF classifier reaches an AUC between 0.62 (cluster 2, Fig. 2C) and 0.55 (cluster 6, FIg. 2D) when trying to distinguish cluster-specific cscGAN-generated cells from real cells, a value that is reasonably close to the perfect situation of random classification (AUC of 0.5) (Table S2).

**Figure 2.**
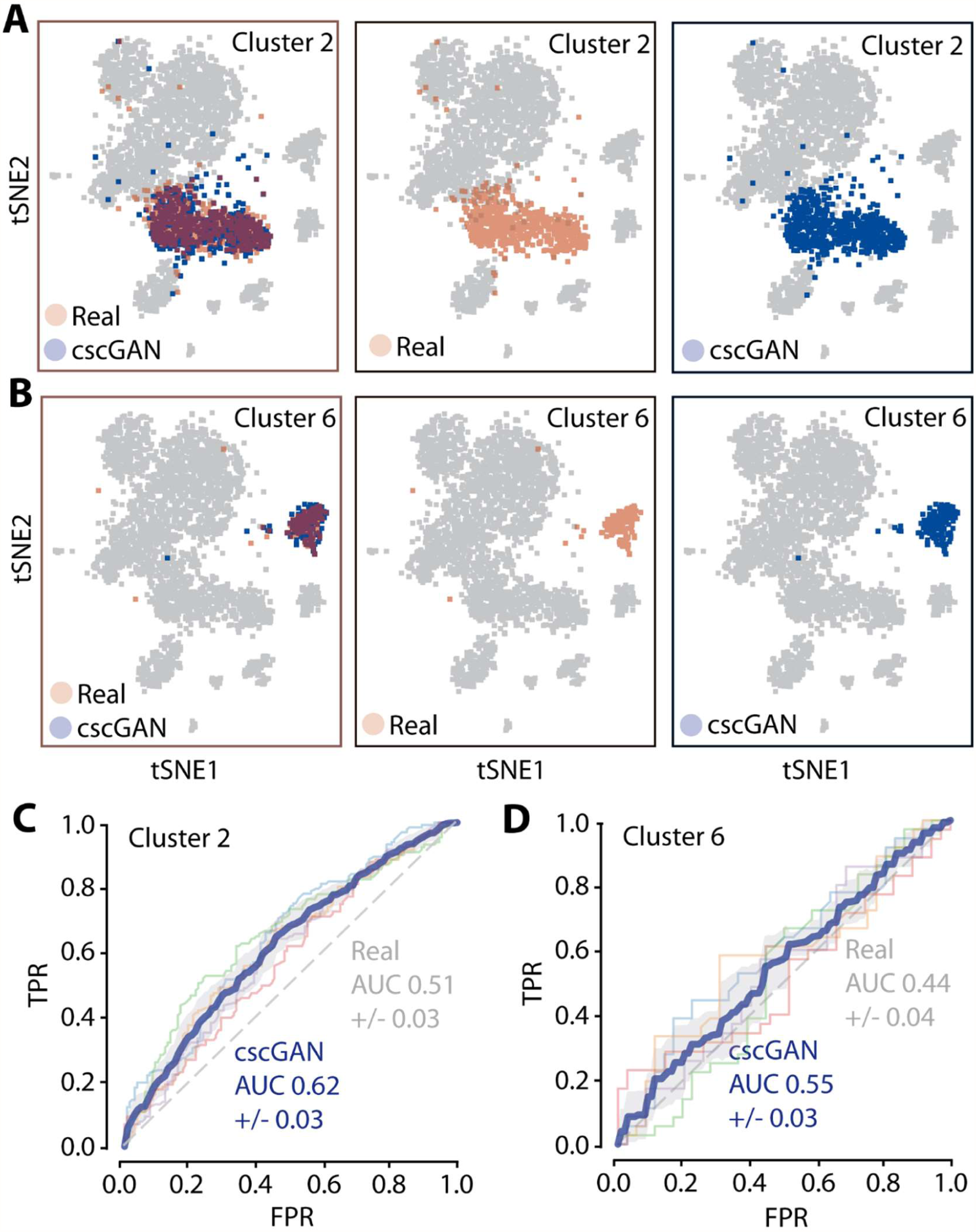
Evaluation of the conditional generation of PBMC cells. A: t-SNE visualization of cluster 2 real cells (red, left and middle panels), cluster 2 generated cells (blue, left and right panels), and other real cells (grey, all panels). B: Same as A for cluster 6 cells. C: Cross-validation ROC curve of a RF classifying cluster 2 real from cscGAN generated cells (cscGAN in blue, chance-level in gray). D: Same as C for cluster 6 cells.

The results of this section demonstrate that the cscGAN can generate high-quality scRNA-seq data for specific clusters or cell types of interest, while rivaling the overall representational power of the scGAN.

### Improved classification of sparse cells using augmented data

We now investigate how we can use the conditional generation of cells to improve the quality and significance of downstream analyses for rare cell populations. The two underlying hypotheses are (i) that a few cells of a specific cluster might not represent the cell population of that cluster well (sampling bias), potentially degrading the quality and robustness of downstream analyses. (ii) This degradation might be mitigated by augmenting the rare population with cells generated by the cscGAN. The base assumption is that the cscGAN might be able to learn good representations for small clusters by using gene expression and correlation information from the whole dataset.

To test the two parts of our hypothesis, we first artificially reduce the number of cells of the PBMC cluster 2 (downsampling) and observe how it affects the the ability of a RF model to accurately distinguish cells from cluster 2 from cells of other clusters. In addition, we train the cscGANs on the same downsampled datasets, generate cells from cluster 2 to augment the downsampled population, retrain an RF with this augmented dataset, and measure the gain in their ability to correctly classify the different populations.

More specifically, cluster 2 comprises 15,008 cells and constitutes the second largest population in the PBMC dataset. Such a large number of cells makes it possible to obtain statistically sound results in various downstream analysis tasks. By deliberately holding out large portions of this population, we can basically quantify how the results would be affected if that population was arbitrarily small. We produce 8 alternate versions of the PBMC dataset, obtained by downsampling the cluster 2 population (keeping 50%, 25%, 10%, 5%, 3%, 2%, 1% and 0.5% of the initial population) (Fig. S8, Table S3). We then proceed to train RF classifiers (for each of those 8 downsampled datasets) (Fig. S9A), on 70% of the total amount of cells and kept aside 30% to test the performance of the classifier (Fig. S9B). The red line in Fig. 3A and Fig. S10 very clearly illustrates how the performance of the RF classifier, measured through the F1 score, gradually but significantly decreases from 0.95 to 0.45 while the downsampling rate goes from 50% to 0.5%.

**Figure 3.**
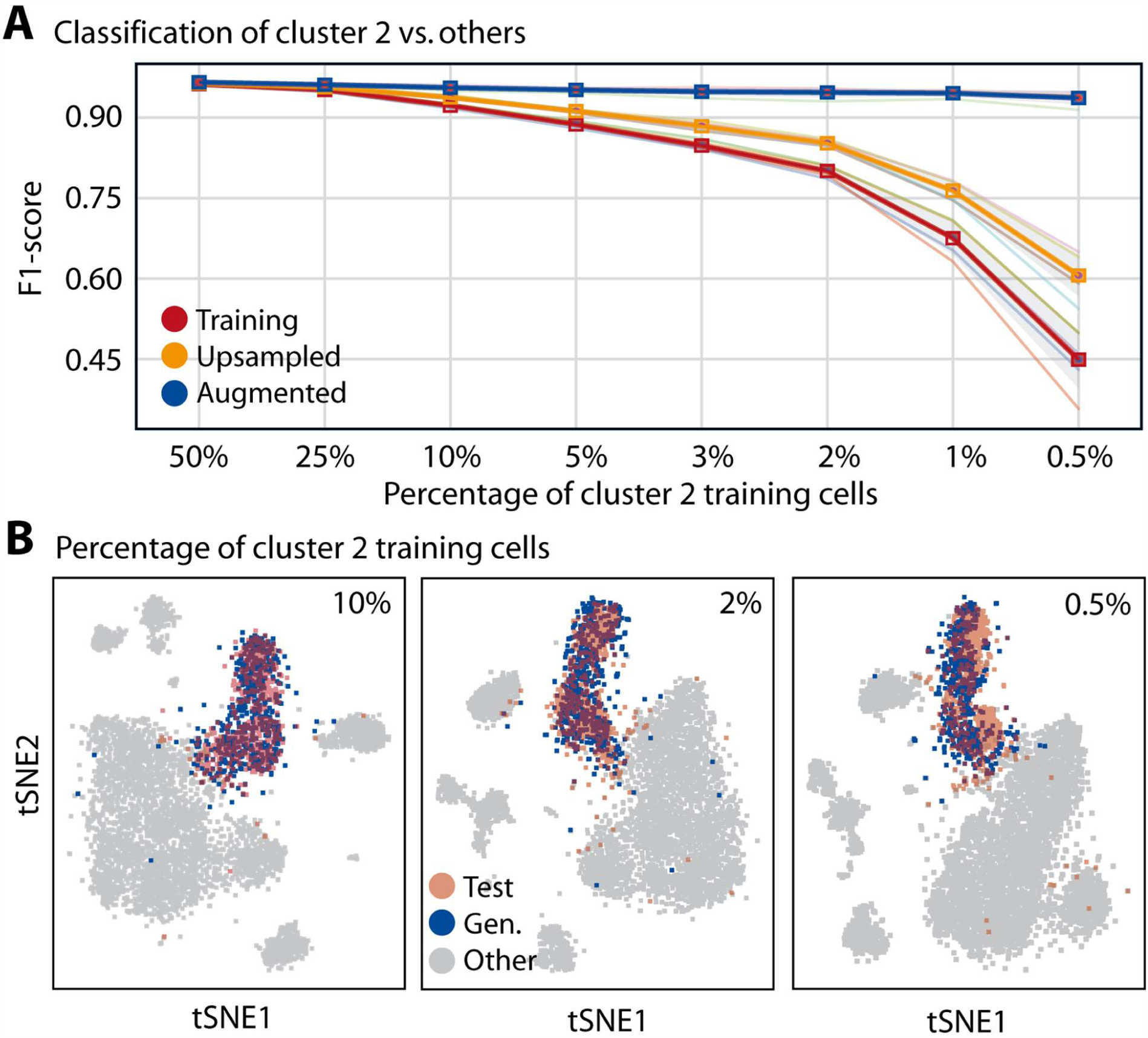
Effects of data downsampling and augmentation on classification and clustering. A: F1-score of a RF classifier trained to discriminate cluster 2 test from other test cells when trained on training (red), upsampled (orange), or augmented (blue) cells for eight levels of downsampling (50% to 0.5%). B: t-SNE representation of cluster 2 real test (red) and cscGAN generated (blue) cells for three levels of downsampling (10%, 2%, and 0.5%). Other trest cells are shown in gray.

To see if we could mitigate this deterioration, we experimented with two ways of augmenting our alternate datasets. First, we used a naïve method, which we call upsampling, where we simply enlarged the cluster 2 population by duplicating the cells that were left after the downsampling procedure (Fig. 9A). The orange line in Fig. 3 shows that this naive strategy actually mitigates the effect of the downsampling, albeit only to a minor extent (F1 score of 0.6 obtained for a downsampling rate of 0.5%). It is important to note that adding noise (e.g. standard Gaussian) to the upsampled cluster 2 cells usually deteriorated the classification performance (data not shown).

In order to understand if *in silico* generated cluster 2 cells could improve the RF performance, we next trained the cscGANs on the 8 downsampled datasets (Fig. S9C). We then proceeded to augment the cluster 2 population with the cells generated by the cscGAN (Fig. 9A). Fig. 3B shows that using as little as 2% (301 cells) of the real cluster 2 data for training the cscGAN suffices to generate cells that overlap with real test cells. When less cells are used the t-SNE overlap of cluster 2 training cells and generated cells slightly decreases (Fig. 3B right panel, Fig. S11). These results strongly suggest that the cluster-specific expression and gene dependencies are learnt by the cscGAN, even when very few cells are available. In line with this assumption, the blue curves in Figure 3A and Fig. S10 show that augmenting the cluster 2 population with cluster 2 cells generated by the cscGAN almost completely mitigates the effect of the downsampling (F1 score of 0.93 obtained for a downsampling rate of 0.5%). Interestingly, the RF improves with increasing numbers of generated cells used for the classifiers’ training (Fig. S12).

Two conclusions can be obtained from these results. First the obvious, few cluster-specific cells do not represent the population well, which we refer to as representational or sampling bias. Second, the usage of cscGAN generated scRNA-seq can significantly mitigate this effect and increases the performance of downstream applications like classification when limited samples of a specific cluster are available.

## Discussion

This work shows how cscGANs can be used to generate realistic scRNA-seq representations of complex scRNA-seq data with multiple distinct cell types and millions of cells. cscGANs outperform current methods in the realistic generation of scRNA-seq data and scale sublinearly in the number of cells. Most importantly, we provide compelling evidence that generating *in silico* scRNA-seq data improves downstream applications, especially when sparse and underrepresented cell populations are augmented by the cscGAN generated cells. We specifically show how the classification of cell types can be improved when the available data is augmented with *in silico* generated cells, leading to classifiers that rival the predictive power of those trained on real data of similar size.

It may be surprising or even suspicious that our cscGAN is able to learn to generate cells coming from very small sub-populations (a few dozens of cells) so well. We speculate that although cells from a specific type may have very specific functions, or exhibit highly singular patterns in the expression of several marker genes, they also share a lot of similarities with the cells from other types, especially with those that share common precursors. In other words, the cscGAN is not only learning the expression patterns of a specific sub-population from the (potentially very few) cells of that population, but also from the (potentially very numerous) cells from other populations. This hypothesis actually aligns with the architecture of cscGAN. In the generator, the only parameters that are cluster specific are those learnt in the Conditional Batch Normalization layers. On the other hand, all the parameters of each of the Fully Connected layers are shared across all the different cell types.

While focusing on the task of cell type classification in this manuscript, many other applications will most probably gain from data augmentation, including - but not limited to - clustering itself, cell type detection, and data denoising. Indeed, a recent manuscript used Wasserstein GANs (WGAN) to denoise scRNA-seq data^21^. For this purpose, the (low-dimensional) representation obtained at the output of the single hidden layer of a critic network was used. These lower-dimensional representations keep cell type-determining factors while they discard ‘noisy’ information such as batch effects. In general, GAN models allow for the simulation of cell development or differentiation through simple arithmetic operations applied in the latent space representation of the GAN, operations for which our conditional cscGANs are especially suited.

It is tempting to speculate how well the scRAN-seq data generation using cscGANs can be applied to other biomedical domains and data types. It is easy to envision, for example, how cscGAN variants could generate realistic (small) RNA-seq or proteomic data. Moreover, cscGAN variants might successfully generate whole genomes with predefined features such as disease state, ethnicity, and sex, building virtual patient cohorts for rare diseases, for example. In biomedical imaging, *in silico* image generation could improve object detection, disease classification, and prognosis, leading to increased robustness and better generalization of the experimental results, extending clinical application.

We hypothesize that data augmentation might be especially useful when dealing with human data, which is notoriously heterogeneous due to genetic and environmental variation. Data generation and augmentation might be most valuable when working with rare diseases or when samples with a specified ethnicity or sex, for example, are simply lacking.

Lastly we would like to emphasize that the generation of realistic *in silico* data has far reaching implications beyond enhancing downstream applications. *In silico* data generation can decrease human and animal experimentation with a concomitant reduction in experimental costs, addressing important ethical and financial questions.

## Methods

### Datasets and pre-processing

#### PBMC

We trained and evaluated all models using a published human dataset of 68,579 peripheral blood mononuclear cells (healthy donor A)^11^. The dataset was chosen as it contains several clearly defined cell populations and is of reasonable size. In other words, it is a relatively large and complex scRNA-seq dataset with very good annotation, ideal for the learning and evaluation of generative models.

The cells were sequenced on Illumina NextSeq 500 High Output with ~20,000 reads per cell. The cell barcodes were filtered as in ^11^ and the filtered genes matrix is publicly available on the 10x Genomics website.

In all our experiments, we removed genes that are expressed in less than 3 cells in the gene matrix, yielding 17,789 genes. We also discarded cells that have less than 10 genes expressed. This, however, did not change the total number of cells. Finally, the cells were normalized for the library-size by first dividing UMI counts by the total UMI counts in each cell and then multiplied by 20,000. See Table S1 for an outlook of this dataset.

#### Brain Large

In addition to the PBMC dataset we trained and evaluated our best performing scGAN model on the currently largest available scRNA-seq dataset of ~1.3 million mouse brain cells (10x Genomics). The dataset was chosen to prove that the model performance scales to millions of scRNA-seq cells, even when the organism, tissue, and the sample complexity varies. The sequenced cells are from the cortex, hippocampus and the subventricular zone of two E18 mice.

The barcodes filtered matrix of gene by cell expression values is available on the 10x genomics website. After removing genes that are expressed in less than 3 cells, we obtained a gene matrix of 22,788 genes. We also discarded cells that have less than 10 genes expressed, which did not affect the overall number of cells. The cells were normalized for the library-size by first dividing UMI counts by the total UMI counts in each cell and subsequent multiplication by 20,000. See Table S1 for an outlook of this dataset.

#### Brain Small

We also examined the performance of the generative models proposed in this manuscript on a subset of the Brain Large dataset provided by 10x Genomics, which consists of 20,000 cells. The pre-processing of the Brain Small dataset was identical to that of the Brain Large dataset, yielding a matrix of 17970 genes by 20.000 cells (Table S1).

#### Louvain clustering

Throughout this manuscript we use the Cell Ranger workflow for the scRNA-seq secondary analysis^11^. First, the cells were normalized by UMI counts. Then, we took the natural logarithm of the UMI counts. Afterwards, each gene was normalized such that the mean expression value for each gene is 0, and the standard deviation is 1. The top 1000 highly variable genes were selected based on their ranked normalized dispersion. PCA was applied on the selected 1000 genes. In order to identify cell clusters, we used Louvain clustering^22^ on the first 50 principal components of the PCA. This replaced the k-means clustering used in Cell Ranger R analysis workflow. We used the Scanpy package^23^ to run the louvain clustering on the first 50 PCs. The number of clusters were controlled by the resolution parameter of scanpy.api.tl.louvain. The higher resolution made it possible to find more and smaller clusters.

For the PBMC dataset we used a resolution of 0.3 which produced 10 clusters. On the other hand, we clustered the Brain Large dataset using a resolution of 0.15 which produces 13 clusters. Finally, the Brain Small dataset was clustered using a resolution of 0.1 which gives 8 clusters.

#### Definition of marker genes

In several experiments, we investigated the expression levels and correlation of genes. For these purposes, a group of 10 ‘marker genes’ was defined by taking the five most highly upregulated genes for the largest two clusters in the dataset (clusters 1 & 2 for the PBMC dataset). Significant upregulation was estimated using the logarithm of the Louvain-clustered dataset and the scanpy.api.tl.rank_genes_groups function with its default parameters (Scanpy 1.2.2)^23^.

### Model description

#### scGAN

In this section, we outline the model used for scGAN by defining the loss function it optimizes, the optimization process, and key elements of the model architecture GANs typically involve two Artificial Neural Networks: a generator, which, given some input random noise, trains to output realistic samples, and a critic that trains to spot the differences between real cells and the ones that the generator produces (Fig. S1). An ‘adversarial’ training procedure allows for those entities to compete against each other in a mutually beneficial way. Formally, GANs minimize a divergence between the distributions of the real samples and of the generated ones. Different divergences are used giving rise to different GAN variants. While original GANs^4^ minimize the so-called Jensen-Shannon divergence, they suffer from known pitfalls making their optimization notoriously difficult to achieve^24^. On the other hand, “Wasserstein GANs” (WGANs)^13,25^ use a Wasserstein distance, with compelling theoretical and empirical arguments.

Let us denote by *P*_*r*_ and *P*_*s*_ the distributions of the real and of the generated cells respectively. The Wasserstein distance between them, also known as the “Earth Mover distance, is defined as follows:

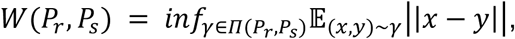

where *x* and *y* are random variables and *Π*(*P*_*r*_, *P*_*s*_)is the set of all joint distributions *γ*(*x,y*)whose marginals are *P*_*r*_ and *P*_*s*_ respectively. Those distributions represent all the ways (called transport plans) you can move “masses” from *x* to *y* in order to transform *P*_*r*_ into *P*_*s*_. The Wasserstein distance is then the cost of the “optimal transport plan”. However, in this formulation, finding a generator that will generate cells coming from a distribution *P*_*s*_ such that it minimizes the Wasserstein distance with the distribution of the real cells, is intractable.

Fortunately, we can use a more amenable, equivalent formulation for the Wasserstein distance, given by the Kantorovich-Rubinstein duality:

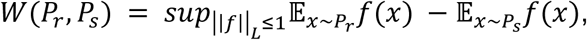

where 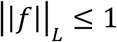 is the set of 1-Lipschitz functions with values in ℝ. The solution to this problem is approximated by training a Neural Network that we previously referred to as the “critic” network, and whose function will be denoted by *ƒ*_*c*_.

The input of the generator are realizations of a multivariate noise whose distribution is denoted by *P*_*n*_. As it is common in the literature, we use a centered Gaussian distribution with unit diagonal covariance (i.e. a multivariate white noise). The dimension of the used Gaussian distribution defines the size of the “latent space” of the GAN. The dimension of that latent space should reflect the “intrinsic dimension” of the scRNA-seq expression data we are learning from, and is expected to be significantly smaller than their “apparent dimension” (i.e. the total number of genes).

If we denote by *ƒ*_*g*_ the function learnt by our generator network, the optimization problem solved by the scGAN is identical to that of a Wasserstein GAN:

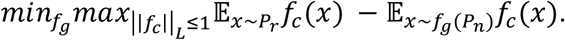

The enforcement of the Lipschitz constraint is implemented using the gradient penalty term proposed by ^25^.

Hence, training a scGAN model involves solving a so-called “minmax” problem. As no analytical solution to this problem can be found, we recourse to numerical optimization schemes. We essentially follow the same recipe as most of the GAN literature^4,24^, with an alternated scheme between maximizing the critic loss (for 5 iterations) and minimizing the generator loss (for 1 iteration). For both the minimization and the maximization, we use a recent algorithm called AMSGrad^26^, which addresses some shortcomings of the widely used Adam algorithm^27^, leading to a more stable training and convergence to more suitable saddle points.

Regarding the architecture of our critic and generator networks, which is summarized in Fig. S1, most of the existing literature on images prescribes the use of Convolutional Neural Networks (CNN). In natural images, spatially close pixels exhibit stronger and more intricate inter-dependencies. Also, the spatial translation of an object in an image usually does not change its meaning. CNNs have been designed to leverage those two properties. However, neither of these properties hold for scRNAseq data, for which the ordering of the genes is mostly arbitrary and fixed for all cells. In other words, there is no reason to believe that CNNs are adequate, which is why scGANs use Fully Connected (FC) layers. We obtained the best results using an MLP with FC layers of 256, 512, and 1024 neurons for the generator and an MLP with FC layrs of 1024, 512, 256 for the critic (Fig. S1B & C). At the outermost layer of the critic network, following the recommendation from^24^, we don’t use any activation function. For every other layer of both the critic and the generator networks, we use a Rectified Linear Unit (ReLU) as an activation function. Naturally, the optimal parameters in each layer of the Artificial Neural Network highly depends on the parameters in the previous and subsequent layers. Those parameters, however, change during the training for each layer, shifting the distribution of subsequent layer’s inputs slowing down the training process. In order to reduce this effect and to speed up the training process, it is common to use Normalization layers such as Batch Normalization^28^ for each training mini-batch. We found that the best results were obtained when using Batch Normalization at each layer of the generator. Finally, as mentioned in the datasets and pre-processing section, each real sample used for training has been normalized for library size. We now introduce a custom Library Size Normalization (LSN) layer that enforces the scGAN to explicitly generate cells with a fixed library size (Fig. S1B).

#### LSN layer

A prominent property of scRNA-seq is the variable range of the genes expression levels across all cells. Most importantly, scRNA-seq data is highly heterogeneous even for cells within the same cell subpopulation. In the field of Machine Learning, training on such data is made easier with the usage of input normalization. Normalizing input yields similarly-ranged feature values that stabilize the gradients. scRNA-seq normalization methods that are used include library-size normalization, where the total number of reads per cell is exactly 20.000 (see also Datasets and pre-processing).

We found that training the scGAN on library-size normalized scRNA-seq data helps the training and enhances the quality of the generated cells in terms of our evaluation criteria (model selection method). Providing library-size normalized cells for training of the scGAN implies that the generated cells should have the same property. Ideally, the model will learn this property inherently. In practice, to speed up the training procedure and make training smoother, we added the aforementioned LSN layer at the output of the generator (Fig. S1B). Our LSN Layer rescales its inputs 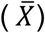 to have a fixed, total read count (*φ*) per cell:

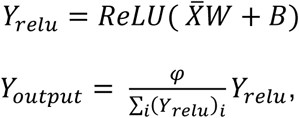

where and *B* are its weights and biases, and (*Y*_*relu*_)_*i*_ denotes the i-th component of the *Y*_*relu*_ vector.

#### cscGAN

Our cscGAN leverages conditional information about each cell type, or sub-population, to enable the further generation of type-specific cells. The integration of such side information in a generative process is known as “conditioning”. Over the last few years, several extensions to GANs have been proposed to allow for such conditioning^19,29,30^. It is worth mentioning that each of those extensions are available regardless of the type of GAN at hand.

We explore two conditioning techniques, auxiliary classifiers (ACGAN)^30^ and projection-based conditioning (PCGAN)^19^. The former adds a classification loss term in the objective. The latter implements an inner product of class labels at the critic’s output. While we also report results obtained with the ACGAN (see Table S2), the best results were obtained while conditioning through projection.

In practice, the PCGAN deviates from the scGAN previously described by: (i) multiple critic output layers, one per cell type and (ii) the use of Conditional Batch Normalization Layers (BNL)^20^, whereby the learnt singular scaling and shifting factors of the BNL are replaced with one per cell type.

As described in Section 2 and 3 of ^19^, the success of the projection strategy relies on the hypothesis that the conditional distributions (with respect to the label) of the data at hand are “simpler”, which helps stabilizing the training of the GAN. When it comes to scRNA-seq data, it is likely that this hypothesis holds as the distribution of the gene expression levels should be “simpler” within specific cell types or sub-populations.

##### Model selection & evaluation

Evaluating the performance of generative models is no trivial task^12^. We designed several metrics to assess the quality of our generated cells at different levels of granularity. We will now describe in detail how those metrics were obtained. They can be grouped into two categories: the metrics we used for model selection (in order to tune the hyper-parameters of our GANs) and the metrics we introduced in the Results section.

#### Metrics used for model selection

As described in the previous section, defining our (c)scGAN model entails carefully tuning several hyper-parameters. We hereby recall the most influential ones: (i) the number and size of layers in the Neural Networks, (ii) the use of LSN layer, and (iii) the use of a Batch Normalization in our generator network.

For each of our models, before starting the training, we randomly pick 3,000 cells from our training data and use them as a reference to measure how it performs. We therefore refer to those 3,000 cells as “real test cells”

To optimize those hyper-parameters, we trained various models and evaluated their performance through a few measures, computed during the training procedure: (a) the distance between the mean expression levels of generated cells and real test cells, (b) the mean sparsity of the generated cells, and (c) the intersection between the most highly variable genes between the generated cells and the real test cells.

First, we compute the mean expression value for each gene in the real test cells. During the training procedure, we also compute the mean expression value for each gene in a set of 3,000 generated cells. The discrepancy is then obtained after computing the Euclidian distance between the mean expression values of the real and the generated cells.

scRNA-seq data typically contains a lot of genes with 0 read counts per cell, which we also use to estimate the similarity of generated and real cells. Naturally, similar sparsity values for real and test cells indicate good model performance whereas big differences indicate bad performance.

Finally, using the Scanpy^23^ package, we estimate the 1,000 most highly variable genes from the real data. During the training, we also estimate what are the 1,000 most highly variable genes from a sample of 3,000 generated cells. We use the size of the intersection between those two sets of 1,000 highly variable genes as a measurement of the quality of the generation.

#### Gene expression and correlation

To highlight the performance of our models, we used violin plots of the expression of several marker genes along with heatmaps displaying the correlation between those same marker genes as expressed among all clusters, or among specific clusters. To produce those plots, we used the expression levels of cells (either test real, or generated by scGAN, cscGAN), in a logarithmic scale. For the heatmaps, we compute the Pearson product-moment correlation coefficients.

#### t-SNE plots

To visualize our generated and real cells within same t-SNE plot, they are embedded simultaneously. In brief, we are trying to assess how “realistic” the generated cells are. Thus our reference point is the real data itself. The delineation of what constitutes noise and what constitutes biologically relevant signal should be driven by the real data only. Hence we project the generated cells on the first 50 principal components that were computed from the real cells in the Cell Ranger pipeline^11^ (see also Datasets and pre-processing). From this low-dimensional representation, we compute the t-SNE embedding.

#### Classification of real vs. generated cells

Building on the 50-dimensional representation of the cells (t-SNE plots section), we trained classifiers to distinguish between real test cells and generated cells. As mentioned in the Results section, we trained Random Forest classifiers with 1000 trees and a Gini impurity quality metric of the decision split using scikit-learn package^31^. The maximum depth of the classifier is set so that, the nodes are expanded until all leaves are pure or until all leaves contain less than 2 samples. The maximum number of features used is the square root of the number of genes.

In order to produce Fig. 1C, which highlights the ability to separate real from generated cells, irrespective of which cluster they are coming from, we used the whole real test set along with generated cells. On the other hand Fig. 2C and 2D are cluster specific (cluster 2 and cluster 5 respectively). We trained the Random Forests using only the cells from those specific clusters. To prevent bias due to class imbalance, each model was trained using an equal number of real test cells and generated cells.

We used a 5-fold cross-validation procedure to estimate how each classifier generalizes. To assess this generalization performance, we plotted the Receiver Operating Characteristic (ROC) curves obtained for each fold, along with the average of all the ROC curves. We also display the AUC in each of those cases. Table S2 reports more extensive results. Each column was obtained for different models, namely cscGAN (based on PCGAN), the conditional scGAN based on ACGAN, while the third column constitutes a baseline obtained when classifying real test cells from real training cells (which virtually constitutes an “impossible” task). Each row represents the performance obtained for cluster specific cells (rows denoted 1 to 6), while the last row (“global”) shows the results obtained for all clusters together.

#### Downsampling

To assess the impact of cluster size on the ability of the cscGAN to model the cluster we artificially reduced the number of cells of the relatively large PBMC cluster 2. We call this approach ‘downsampling’ throughout the manuscript.

Eight different percentages {50%, 25%, 10%, 5%, 3%, 2%, 1%, 0.5%} of cluster 2 cells were sampled using a random seed and a uniform sampling distribution over all the cluster 2 cells (Table S3). We sampled nested subsets (for each seed, the smaller percentage samples are a complete subset of the larger ones). In order to accurately estimate the generalization error across the different experiments and to avoid potential downsampling (sampling bias) artifacts, we conducted all our experiments using five different random seeds. For the classification of cell subpopulations (see next paragraph) we report the average as well as the 5 individual values for the different seeds.

#### Classification of cell sub-populations

To investigate the use of the proposed cscGAN model for data augmentation, we examined the performance of cell subpopulation classification before and after augmenting the real cells with generated cells. For this purpose, and as described in the previous paragraph and the Results section, we produced alternate datasets with sub-sampled cluster 2 populations (Fig. S8).

For simplicity, we focus in this section on the experiment where cluster 2 cells were downsampled to 10% using five different random seeds (Fig. S9). We advise to use Figure S9 as an accompanying visual guide to this text description.

Using the previously introduced 50-dimensional PCA representation of the cells, three RF models were trained to distinguish cluster 2 cells from all other cell populations (**RF downsampled, RF upsampled**, and **RF augmented**) (Fig. S9A). In the training data for all the three classifiers, 70% of the cells from all the clusters except cluster 2 (i.e. 37,500 cells) were used (light blue boxes in Fig. S9A).

##### RF downsampled

For the first RF classifier, we used 10% of cluster 2 cells (1,502 cells) and 70% other cells (37,500 cells) to train the RF model. We refer to this dataset as the “RF downsampled” set in Fig. S9A. This dataset was also used to train the cscGAN model, which is used later to generate *in silico* cluster 2 cells (Fig. S9C). It is important to note that RF classifiers for ‘RF downsampled’ datasets always use weights that account for the cluster-size imbalance. The reason for this is that RFs are sensitive to unbalanced classes, leading to classifiers that always predict the much larger class, thereby ‘optimizing’ the classification error^32^.

##### RF upsampled

For the second RF classifier, we uniformly sampled with replacement 5,000 cells from the 1,502 cluster 2 cells (10%). We added those 5,000 (copied) cells to the original 1,502 cells. We refer to this dataset as “RF upsampled” in Fig. S9A. Again, the rationale for this ‘upsampling’ is that RF classifiers are sensitive to class imbalances, as outlined in the previous paragraph (RF downsampled). Of note, we have also conducted experiments where we added standard Gaussian noise to the upsampled cells, which always reduced the performance of the RF classifier and are therefore not shown.

##### RF augmented

Finally, the third classifier training data “RF augmented” consists of 10% cluster 2 cells as well as 5,000 cluster 2 cells generated using the 10% cscGAN model as shown in Fig. S9C. The 10% cscGAN model was trained on 10% cluster 2 cells as well as all other cells (53,571 cells, Fig. S9C).

The RF classifiers were trained using the same parameters as descirbed in the ‘Generated vs. real classification’ methods section, using 1,000 trees and ‘Gini impurity’. The only difference is that here the class weights during the training are adjusted inversely proportional to the class frequencies. This approach of adjusting class weights is of paramount importance when an imbalanced training data is used^32^. The scikit-learn package^31^ was used to conduct all experiments to classify cell sub-populations.

Test cells consisted of 30% of the data from all the clusters. Since we are testing the cscGAN’s ability to augment different percentages of real cluster 2 cells, we made sure that the 30% of cluster 2 cells used in the test set were selected from the cells which were not seen by any trained cscGAN model (Fig S8 & S9B).

To prove that the downsampling limits the ability to classify and that augmenting the dataset mitigates this effect, all three RF classifiers were trained to classify cluster 2 cells vs. all other sub-populations. The F1 score of each classifier is calculated and presented in different colors (Fig. 3A).

Furthermore, in order to understand how augmentation helps to separate close clusters, we trained the same three RF classifiers after removing all clusters except cluster 2 and 1 from the corresponding training data. We repeated this procedure for cluster 2 and 5, and cluster 2 and 3. We chose those clusters in particular because their highly differentially expressed genes are also highly expressed in 2 meaning that separating them from cluster 2 is more difficult (Fig. 1A). In a similar way, F1 scores for classification of cluster 2 vs. 1 (Fig. S10A), and cluster 2 vs. 3 (Fig. S10B), and cluster 2 vs. 5 (Fig. S10C) are calculated and reported.

As mentioned above, we repeated this procedure for different downsampling levels of cluster 2 cells and for 5 different sampling seeds for each level (Table S3).

When training the RF classifier with the augmented dataset, the number of cells used in the augmentation was set to 5,000 cells. This, however, doesn’t necessarily mean that 5,000 cells is the optimal number of cells to be added. The increase in the F1 score due to augmenting the data with generated cells depends on two factors: (i) the number of real cells in the original sub-population and (ii) the number of cells used for augmentation. To highlight the impact of the number of generated cells used for data augmentation, we trained the previously mentioned RF classifiers using different numbers of generated cells (from 100 to 12,000) while keeping the number of other cells constant (see Fig. S11).

#### Splatter comparison

In addition to what has been previously introduced in the Results section, we also compared the performance of the scGAN to Splatter^16^, using the metrics described in their manuscript. Briefly, Splatter simulation is based on a gamma-Poisson hierarchical model, where the mean expression of each gene is simulated from a gamma distribution and cell counts from a poisson distribution. We noticed that Splatter uses the Shapiro– Wilk test to evaluate the library size distribution, which limits the number of input cells to 5,000. Therefore, we slightly modified the code that allows Splatter to take more than 5,000 cells as input.

While scGAN learns from and generates library-size normalized cells, Splatter is not suited for that task. For the sake of fairness, we used the Splatter package on the non-normalized PBMC training dataset. We then generated (non-normalized) cells, which we normalized, so that they could be compared to the cells generated by scGAN. Following ^16^, we used the following evaluation metrics: distribution of the mean expression, of the variance, of the library sizes and ratio of zero read counts in the gene matrix. The results were computed using the Splatter package and are reported in Fig. S5.

We observe that the results obtained by Splatter are marginally better than or identical to those of the scGAN (Fig. S3). The results from those measures suggest that both Splatter and scGAN constitute almost perfect simulations. However, Splatter simulates “virtual” genes. While those genes share some characteristics with the real genes Splatter infers its parameters from, there is no one-to-one correspondence between any “virtual gene” simulated in Splatter generated cells and the real genes. It is therefore non-sensical to compare Splatter-simulated cells with real cells, as we did to evaluate the quality of (c)scGAN-generated cells. This also prohibits the use of Splatter for data-augmentation purposes.

This being said, we also would like to pinpoint that while the (c)scGAN is able to capture the gene-gene dependencies expressed in the real data (Fig. 1B and S7), this does not hold for Splatter, for which the “virtual” genes are mostly independent from each other. To prove this point, we extract the 100 most highly variable genes from the real cells, the cells generated by Splatter, and the cells generated by the scGAN. We then proceed to compute the Pearson correlation coefficients between each pair within those 100 genes (Fig. S4). It reveals that while those most highly variable genes in the real cells or those generated by the scGAN exhibit some strong correlations, highly variable genes are mostly independent from each other in the cells generated by Splatter. As already mentioned in the Results section, the co-regulation of genes is an important aspect of biological gene-regulatory networks and should be taken into account when generating *in silico* data for algorithmic testing.

##### Software, packages, and hardware used

Our (c)scGAN Tensorflow^33^ implementation can be found on https://github.com/imsb-uke/scGAN, including documentation for the training of the (c)scGAN models. As mentioned several times, we used Scanpy^23^ to conduct most of the data analysis. We also compared our results to those of Splatter^16^, and adapted the code they provided on Github (https://github.com/Oshlack/splatter).

For the sake of reproducibility, here is a list of the version of all the packages we used:

- Tensorflow: 1.8
- Scanpy: 1.2.2
- Anndata: 0.6.5
- Pandas: 0.22.0
- Numpy: 1.14.3
- Scipy: 1.1.0
- Scikit-learn: 0.19.1
- R: 3.5.0 (2018-04-23)
- loomR 0.2.0
- SingleCellexperiment 1.2.0
- Splatter 1.4.0

## List of abbreviations

scGAN: single-cell Generative Adversarial Network

cscGAN: conditional single-cell Generative Adversarial Network

RNA-seq: RiboNucleic Acid sequencing

scRNA-seq: single-cell RiboNucleic Acid sequencing

GAN: Generative Adversarial Network

PBMC: Peripheral Blood MonoCytes

t-SNE: t-distributed Stochastic Neighbor Embedding

RF: Random Forest

ROC: Receiver Operating Characteristic

AUC: Area Under the (ROC) Curve

GPU: Graphics Processing Unit

LSN: Library Size Normalization

UMI: Unique Molecular Identifiers

PCA: Principal Component Analysis

PCs: Principal Components

CNN: Convolutional Neural Network

FC: Fully-Connected

ReLU: Rectified Linear Unit

ACGAN: Auxiliary Classifier Generative Adversarial Network

## Contributions

SB initiated the project. SB, PM, MM, and DSM designed the study, deep learning models, and analysis. PM, MM, and DSM built the deep learning models. PM, MM, VB, and CK analyzed the data. SB, PM, MM, DSM, and VB contributed to the manuscript.

## Supplementary Figures & Tables

**Table S1.**
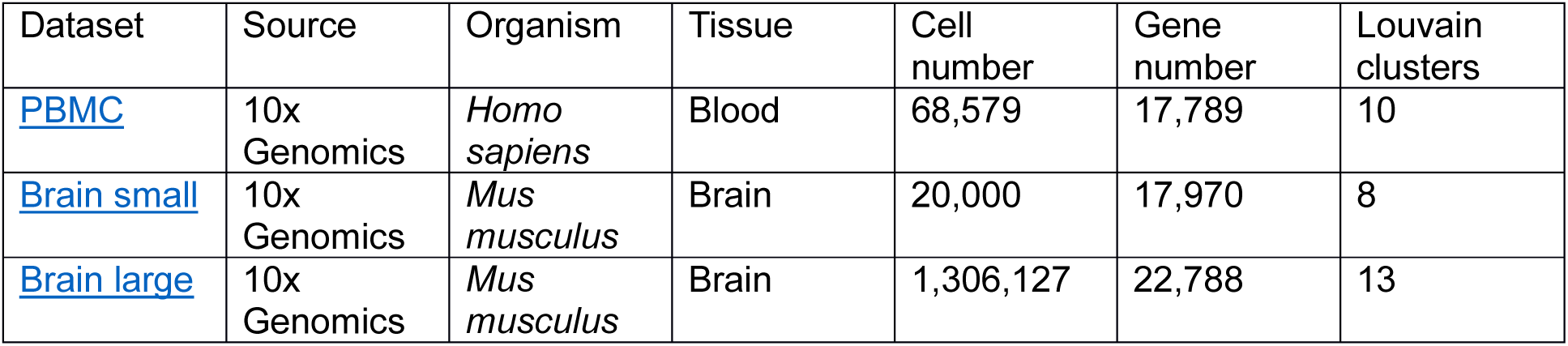
scRNA-seq datasets used. Description of the datasets used throughout the manuscripts, displaying the species and tissue of origin, the number of cells and genes expressed, and the number of clusters inferred with the Louvain method.

**Figure S1.**
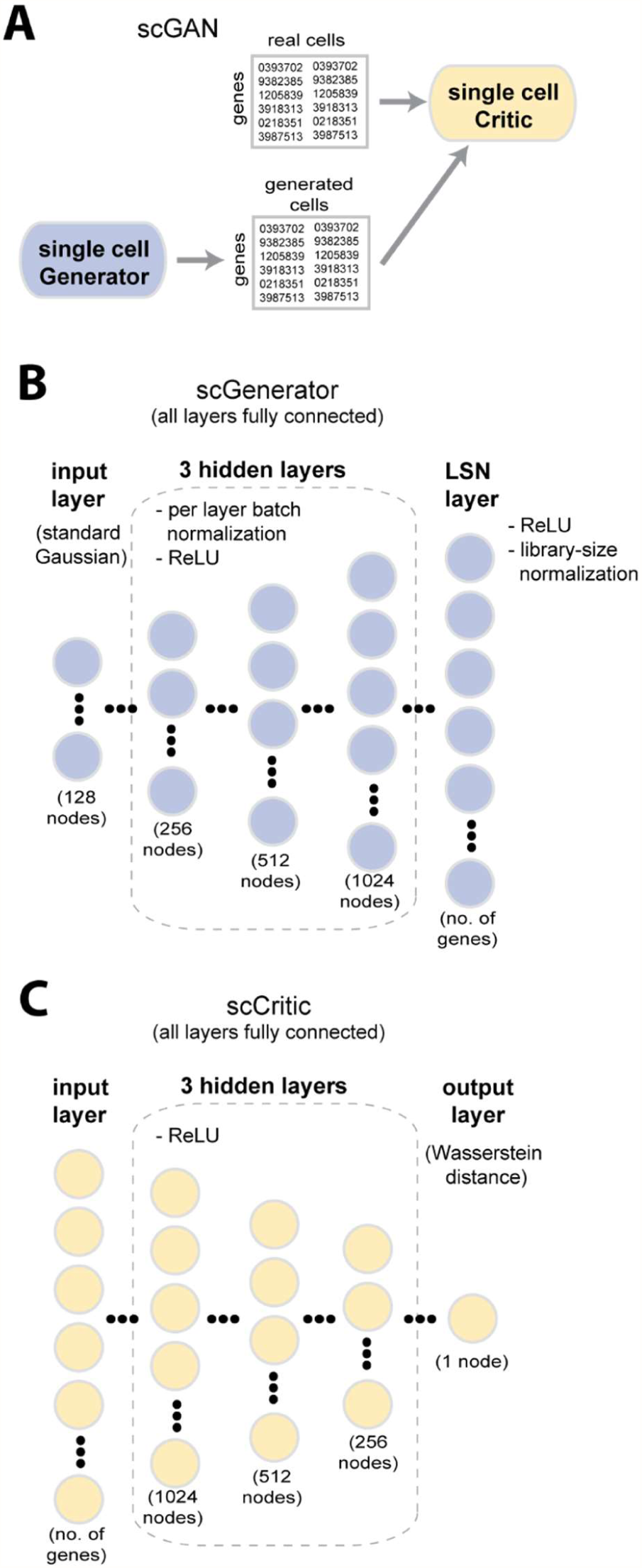
Schematic representation of the scGAN. A: High-level architecture of the scGAN. B: Architecture of the generator network. The generator consists of a Fully-Connected network with three hidden layers of growing size, each featuring Batch Normalization and ReLU activation, and a Library-Size Normalization output layer. The inputs are realizations of standard Gaussian noise. C: Architecture of the critic network. The critic consists of a Fully-Connected network with three hidden layers of decreasing size with ReLU activation.

**Figure S2.**
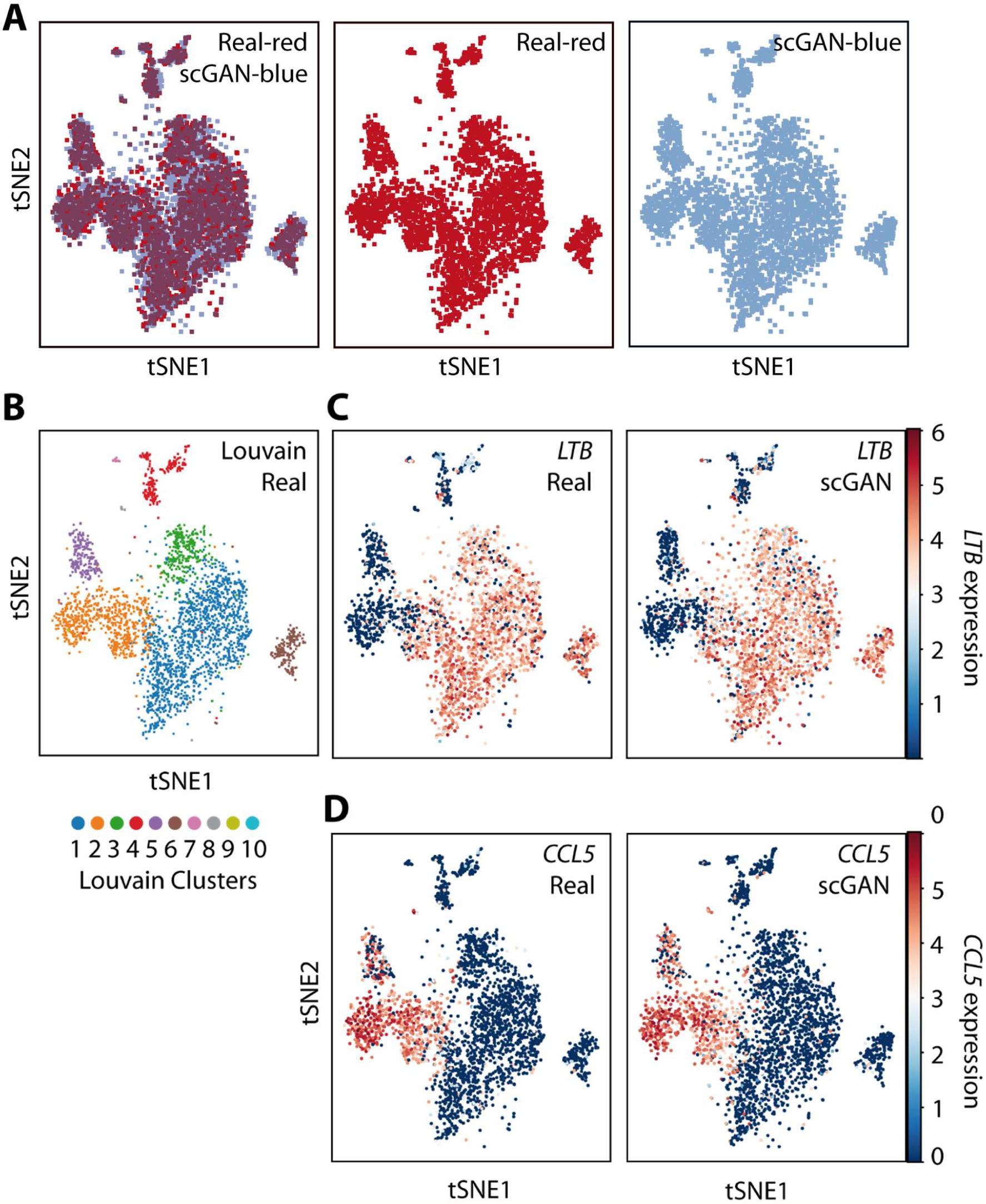
t-SNE visualizations of real and scGAN generated PBMC cells. A: Real cells are shown in red (left and middle panel) and generated cells in blue (left and right panels). B: Real cells are shown with their Louvain clustering. C: LTB gene expression for real (left) and scGAN generated (right) cells. D: CCL5 gene expression for real (left) and scGAN generated (right) cells.

**Figure S3.**
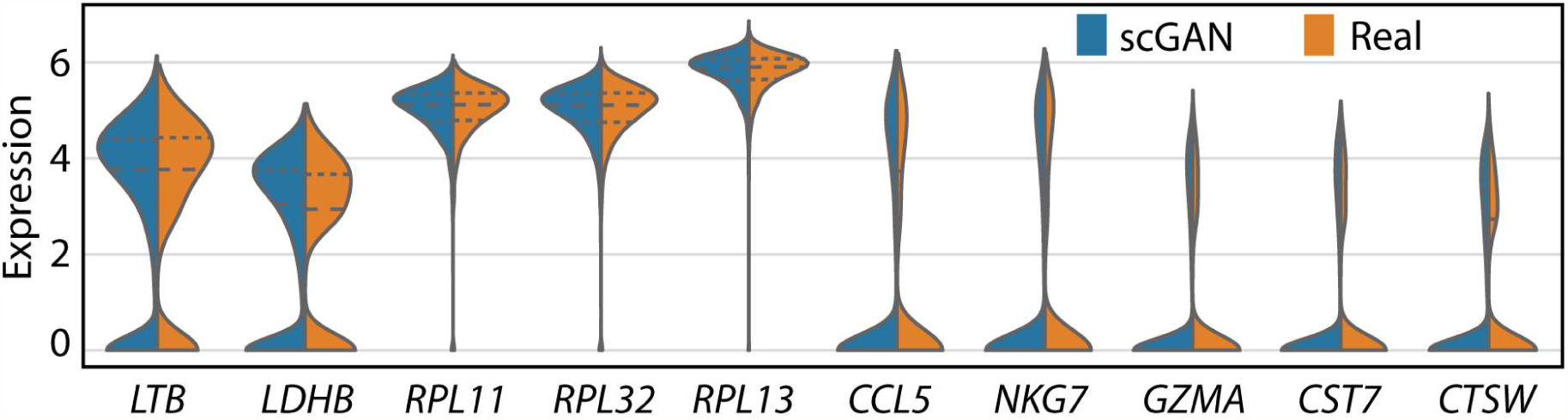
Expression of ten marker genes for real and generated cells. Split violin plots of the distribution of the top five marker genes of cluster 1 (LTB, LDHB, RPL11, RPL32, RPL13) and cluster 2 (CCL5, NKG7, GZMA, CST7, CTSW). Blue corresponds to scGAN generated cells, orange to real data.

**Figure S4.**
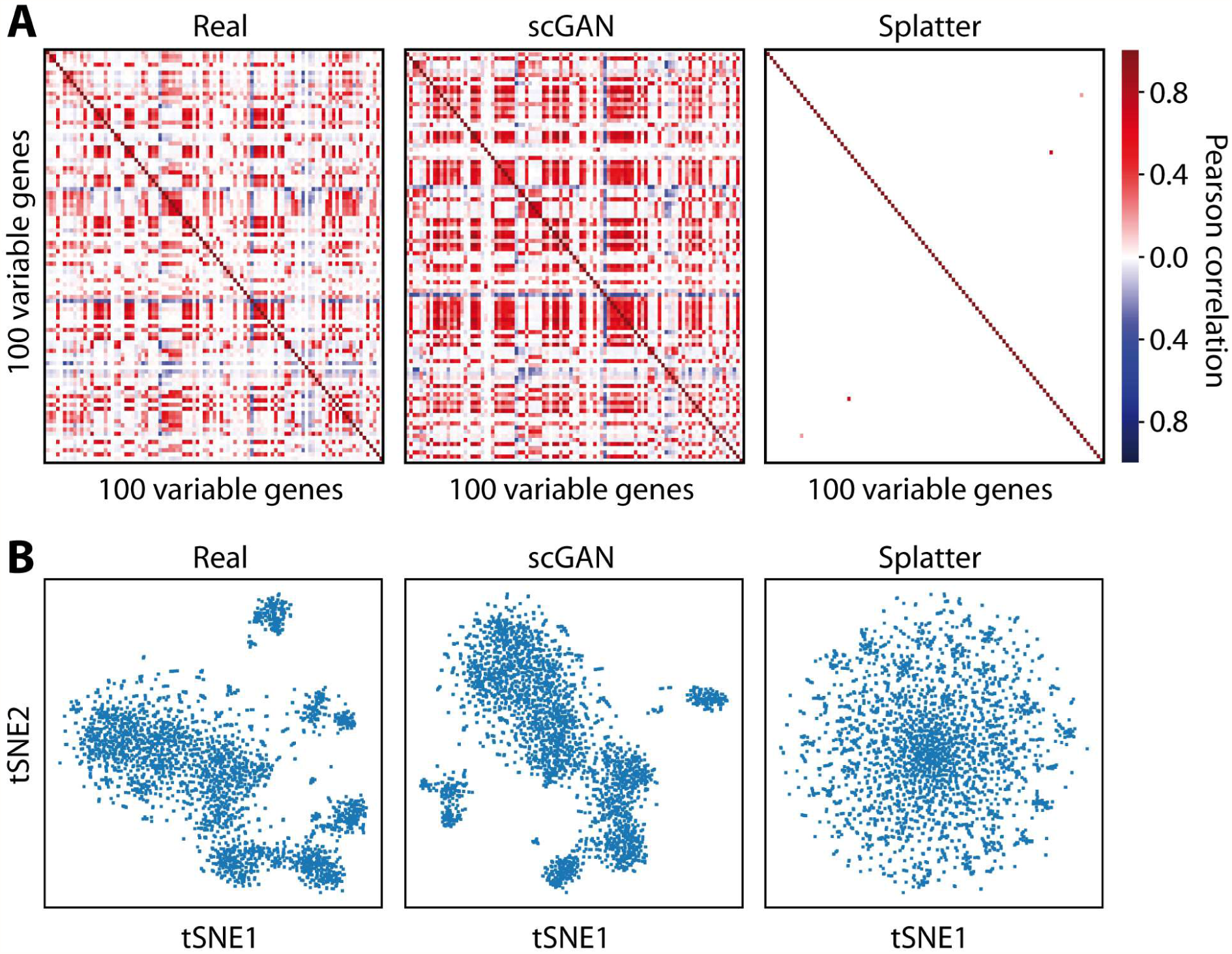
Comparison of Real, scGAN-generated, and Splatter-generated gene correlations and cell clustering. A: Pearson correlation of the 100 most highly variable genes for Real (left), scGAN-generated (middle), and Splatter-generated (right) data. It should be noted that the 100 most highly variable genes were calculated for Real, scGAN, and Splatter data separately, as Splatter does not keep the gene information of the original data. Model parameters were learnt using the PBMC data. B: t-SNE visualizations of Real (left), scGAN-generated (middle), and Splatter-generated (right) cells. It is to be noted that different t-SNE embeddings were used for each t-SNE plot since Splatter does not keep the gene information of the original data. Models were learnt using the PBMC data.

**Figure S5.**
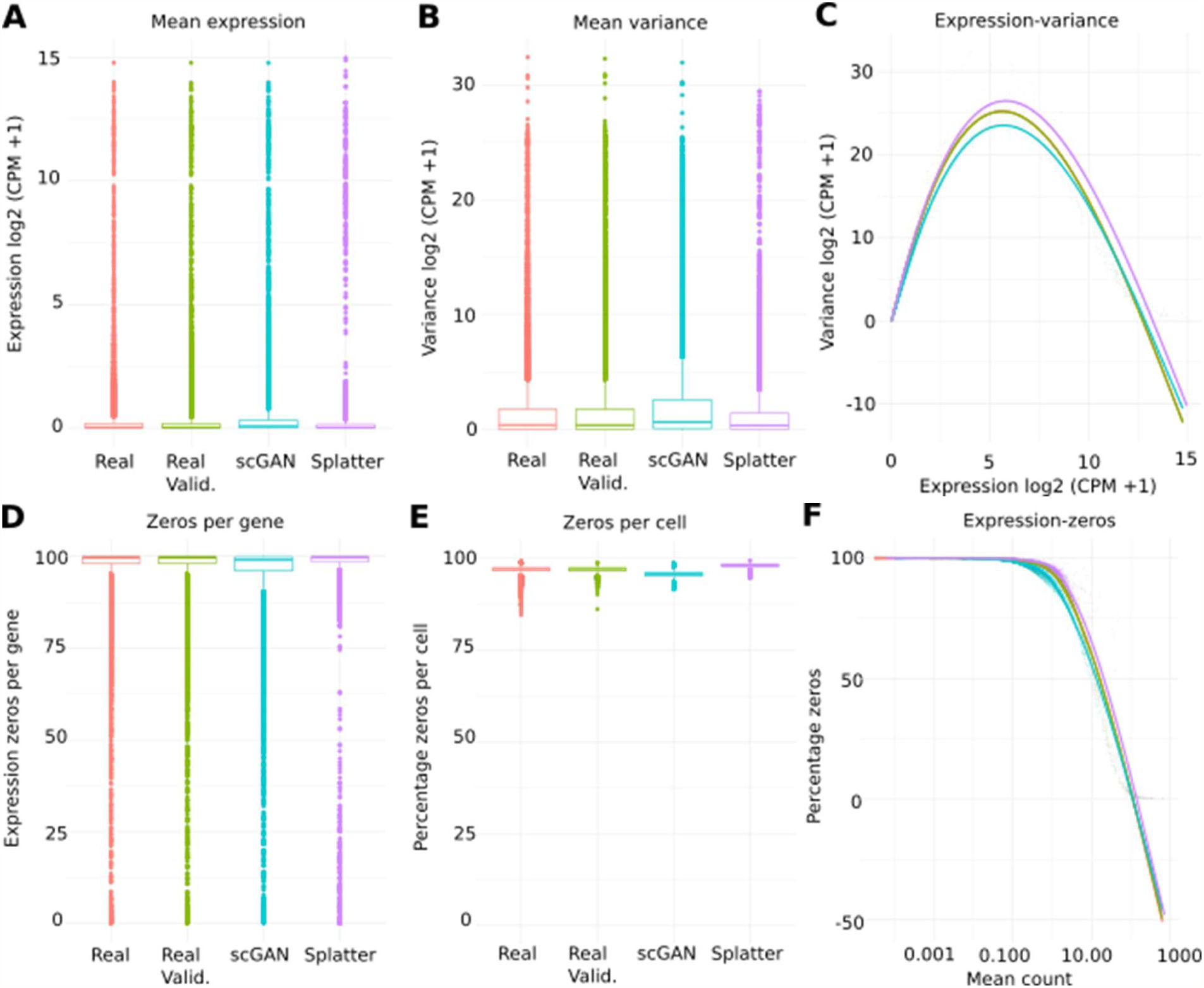
Basic statistical evaluation of the real and scGAN and Splatter generated data. A: Box plot of the mean expression per cell in real training (red), real test (green), scGAN generated (turquoise), and Splatter generated (purple) cells. B: Box plot of the mean variance per cell. C: Mean variance (y-axis) against the per cell mean expression (x-axis). D: Box plot of the percentage of zero expression values per gene. E: Box plot of the percentage of zero expression values per cell. F: Mean count (x-axis) against the percentage of zero expressed genes per cell (y-axis).

**Figure S6.**
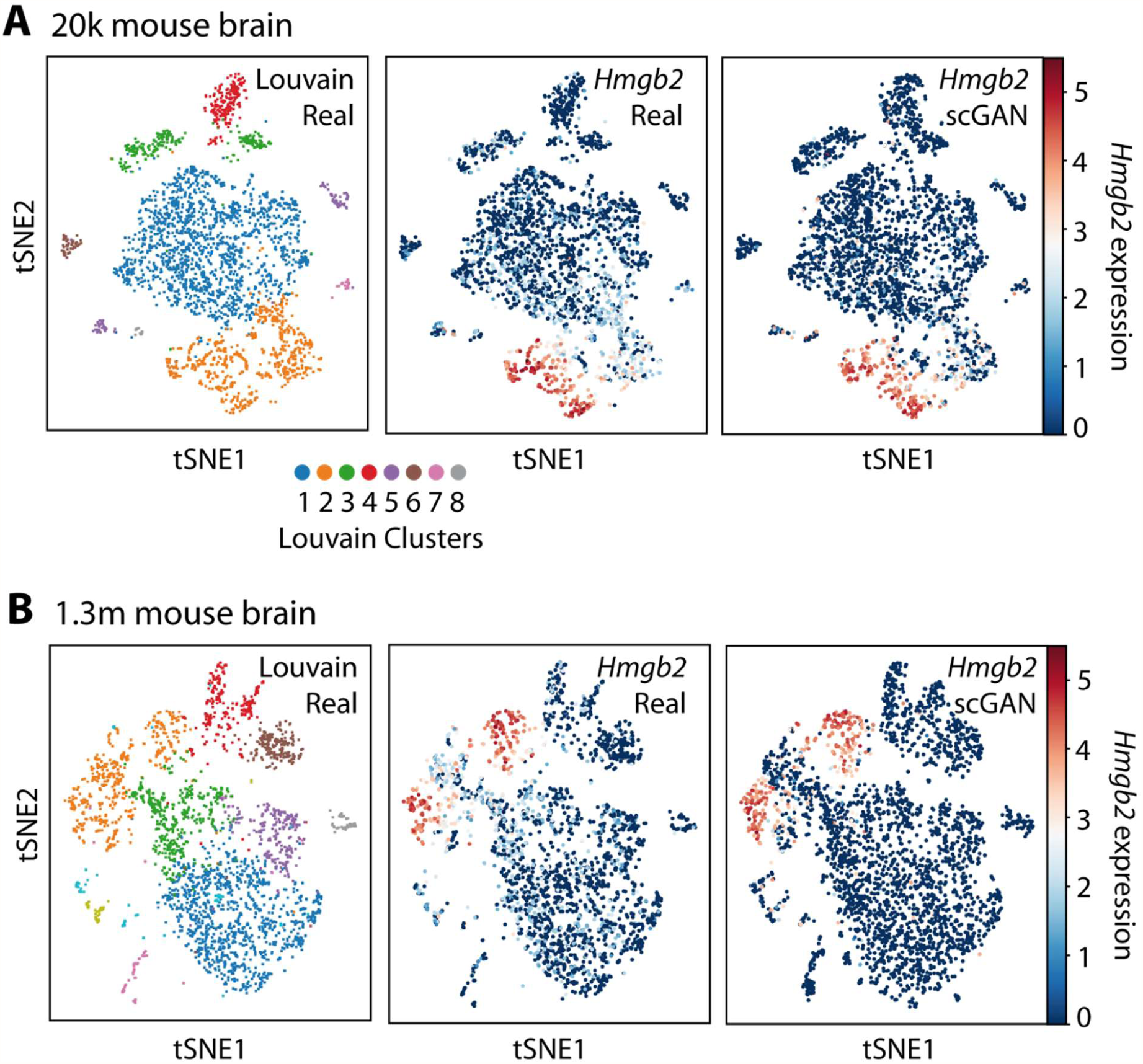
Evaluation of the scGAN simulated Brain Small and Brain Large cells. A: t-SNE visualization of Louvain-clustered real cells (left) and the Hmgb2 gene expression in real (middle) and scGAN generated (right) cells for the Brain Small dataset (20k mouse brain). B: t-SNE visualization of Louvain-clustered real cells (left) and the Hmgb2 gene expression in real (middle) and scGAN generated (right) cells for the Brain Large dataset (1,3 million mouse brain).

**Figure S7.**
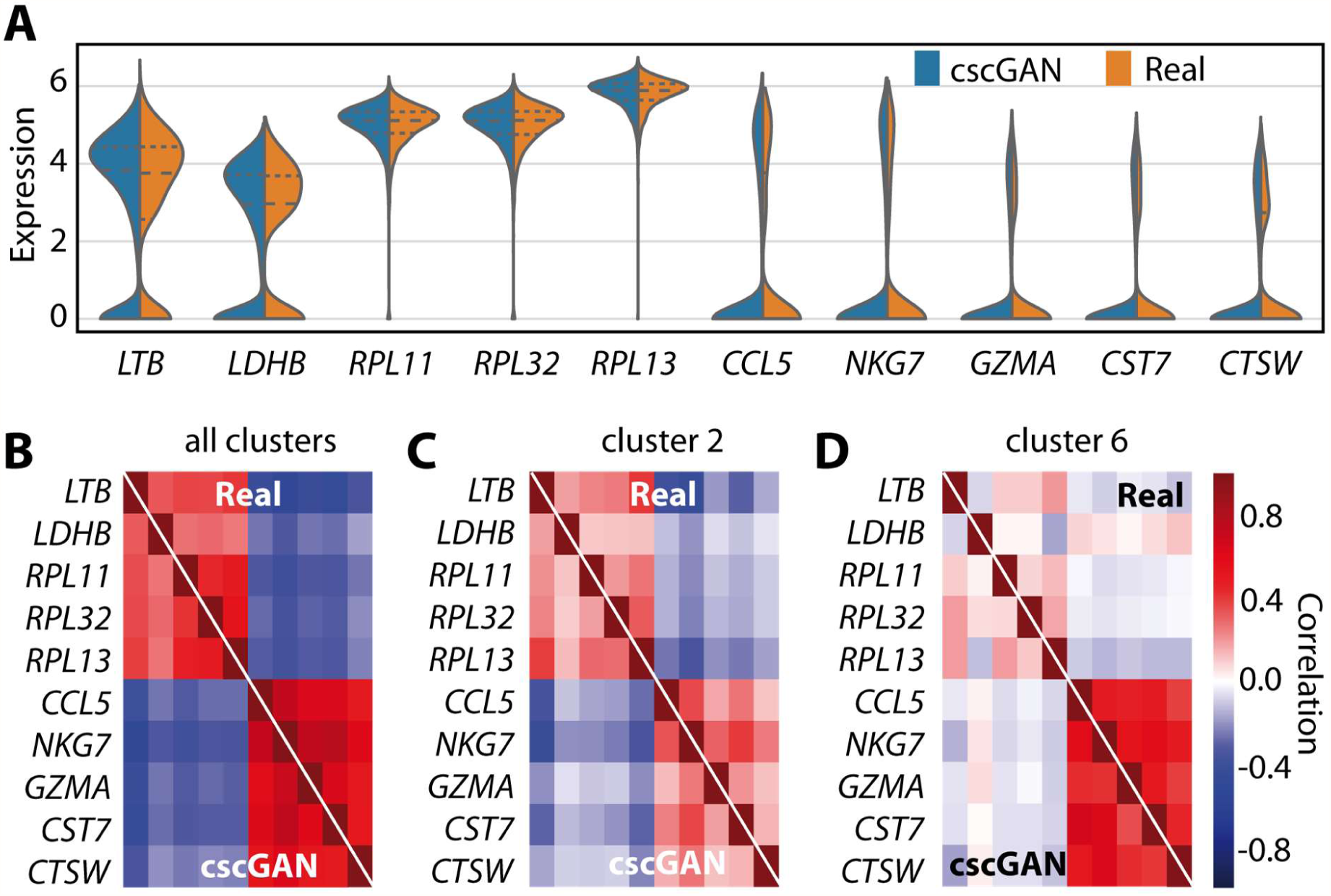
Expression and correlation of ten marker genes for real and conditionally generated PBMC cells. A: Split violin plots of the distribution of the top five marker genes of cluster 1 (LTB, LDHB, RPL11, RPL32, RPL13) and cluster 2 (CCL5, NKG7, GZMA, CST7, CTSW). Blue corresponds to cscGAN generated cells, orange to real data. B-D: Pearson correlation of marker genes for the scGAN generated (bottom left) and the real (upper right) data for (B) all cells, (C) cluster 2, and (D) cluster 6 cells.

**Table S2.** Overview of RF classification performance discriminating real from cscGAN generated cells. Cross-validation area under the ROC curve (AUC) of RFs classifying between real and cscGAN generated cells using a projection (Projection GAN) or an ACGAN critic. As control we also show the classification performance on real training data, which should have chance-level performance (Real). The first six rows correspond to the classification performance for specific clusters (clusters 1-6, other clusters are to small for proper classification), while the last row highlights the classification performance across all clusters (clusters 1-10). For each cell of this table, the left value represents the average AUC across the five folds of the cross-validation. The right value corresponds to the standard deviation.

| Cluster | Projection GAN | ACGAN | Real |
| --- | --- | --- | --- |
| 1 | 0.65 ± 0.02 | 0.76 ± 0.02 | 0.55 ± 0.01 |
| 2 | 0.62 ± 0.03 | 0.69 ± 0.01 | 0.51 ± 0.03 |
| 3 | 0.61 ± 0.05 | 0.78 ± 0.01 | 0.55 ± 0.06 |
| 4 | 0.60 ± 0.07 | 0.98 ± 0.01 | 0.51 ± 0.03 |
| 5 | 0.63 ± 0.07 | 0.66 ± 0.08 | 0.44 ± 0.03 |
| 6 | 0.55 ± 0.03 | 0.98 ± 0.02 | 0.44 ± 0.04 |
| All | 0.63 ± 0.01 | 0.69 ± 0.01 | 0.53 ± 0.01 |

**Figure S8.**
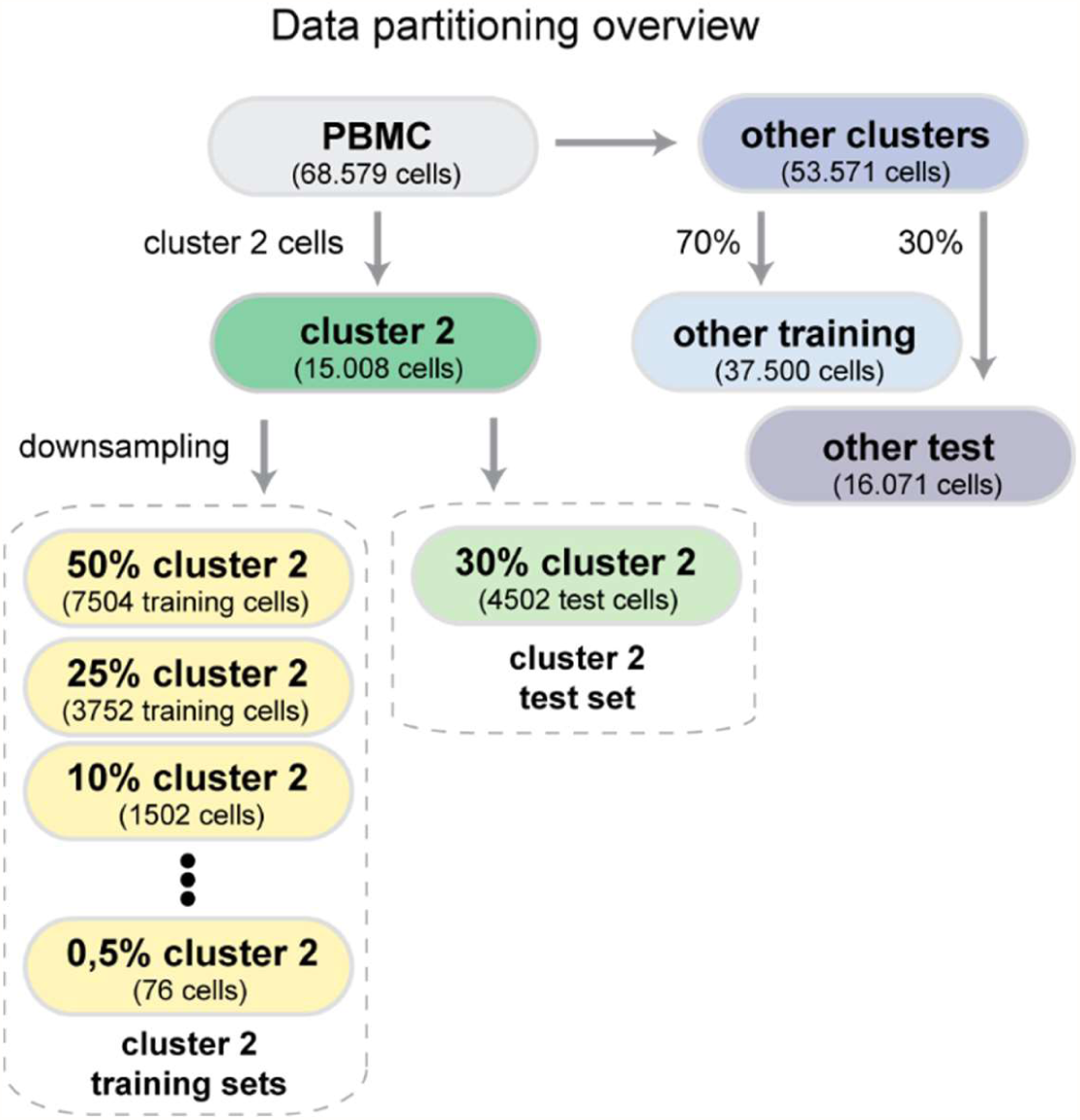
Overview of the PBMC data partitioning for downsampling and augmentation experiments. Cells from the cluster 2 population (dark green) are split into a cluster 2 training set that is downsampled into eight datasets with different cell numbers (yellow), and a test set of 30% of all cluster 2 cells (light green). Cells from other clusters are split into a training set (other training, light blue, 70% of other clusters set) and a test set (other test, dark blue, 30% of other clusters set).

**Figure S9.**
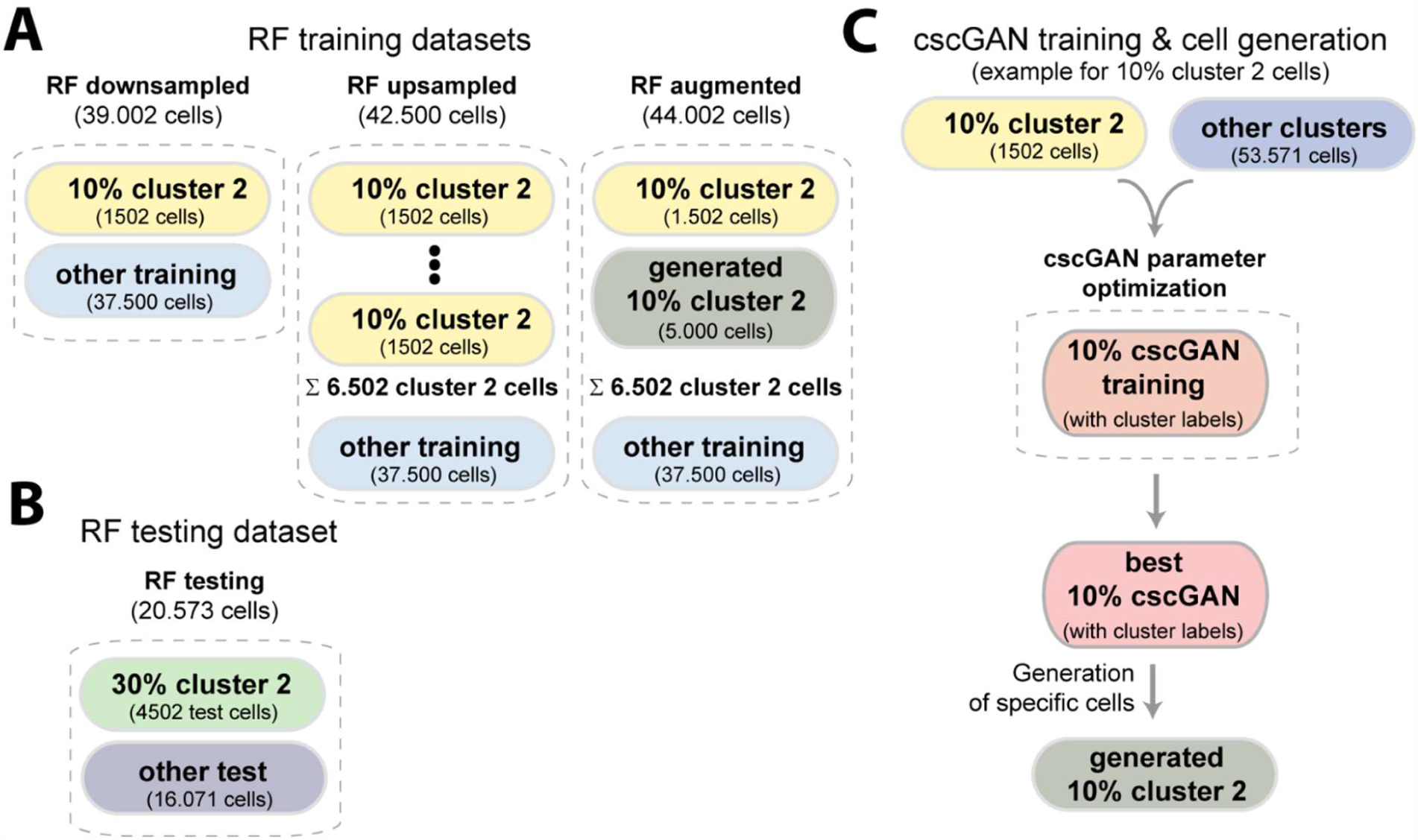
Schematic representation of the datasets used for classification and cell generation. A: RF training was conducted on three different datasets for each percentage of downsampling of cluster 2 cells (as an example we use 10% in the figure). The RF downsampled dataset consists of the 10% cluster 2 set (yellow). The RF upsampled dataset contains 6,502 cluster 2 cells sampled with replacement from the 10% cluster 2 set. The RF augmented dataset contains the 10% cluster 2 set, and 5,000 cells generated using the generated 10% cluster 2 set (grey, see also panel C). In addition, all three datasets contain the other training set (light blue). B: RF testing was conducted on the 30% cluster 2 test set (green) and the other test set (dark blue). C: For data augmentation (see RF augmented) we generate cluster 2 cells using a cscGAN. The cscGAN is trained on the 10% cluster 2 set (yellow) and the other clusters set (blue), yielding a generated 10% cluster 2 set (gray).

**Table S3.**
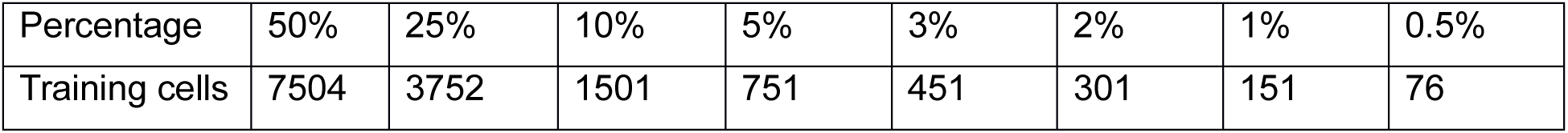
*Number of cluster 2 cells used for all eight levels of downsampling*.

| Percentage | 50% | 25% | 10% | 5% | 3% | 2% | 1% | 0.5% |
| --- | --- | --- | --- | --- | --- | --- | --- | --- |
| Training cells | 7504 | 3752 | 1501 | 751 | 451 | 301 | 151 | 76 |

**Figure S10.**
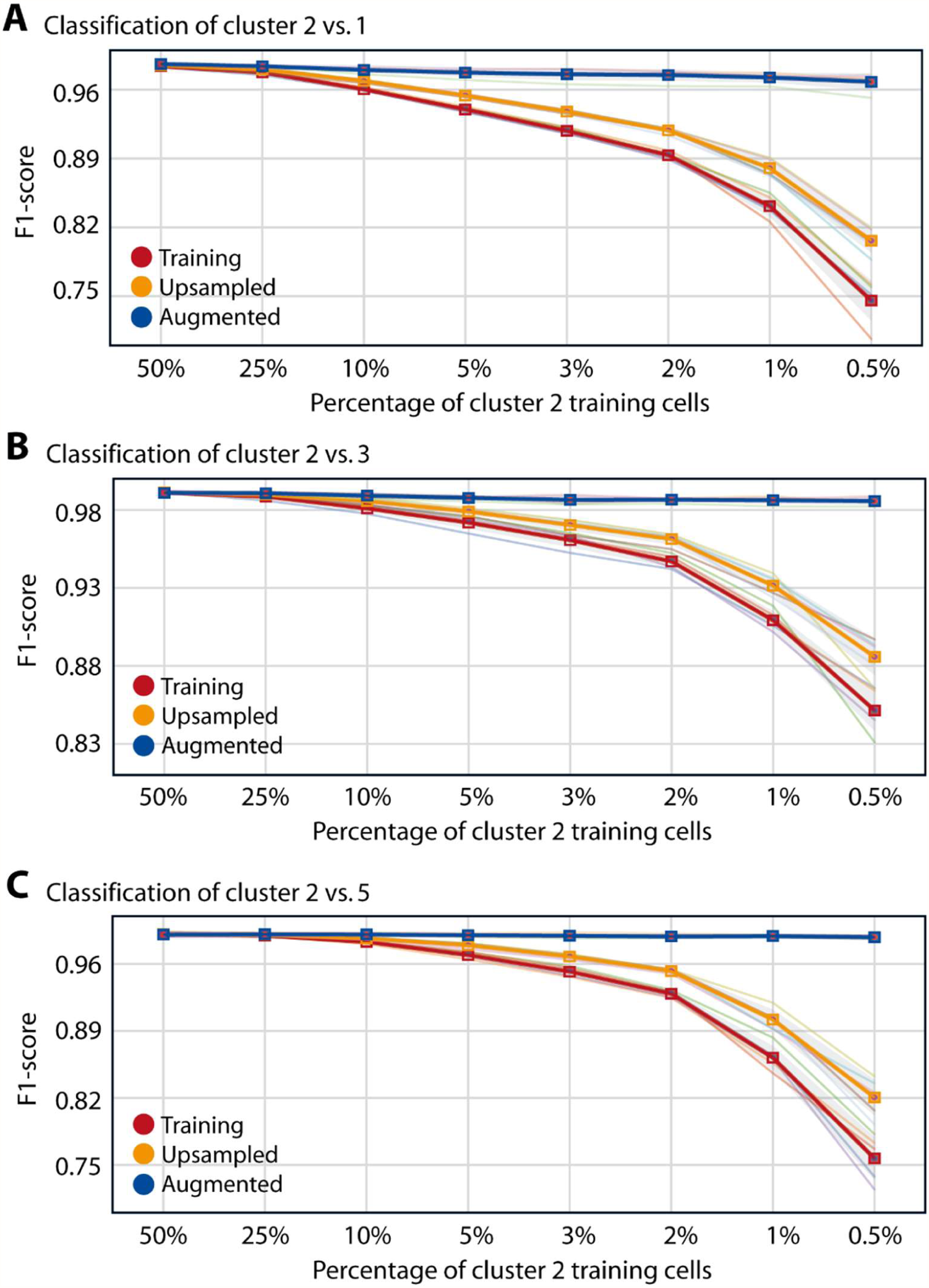
RF classification performance for three cluster-specific comparisons. F1-score reached by a RF classifier trained to discriminate (A) cluster 2 from cluster 1, (B) cluster 2 from cluster 3, and (C) cluster 2 from cluster 5 cells. The RFs were trained on training (red), upsampled (yellow), or augmented (blue) datasets for eight different levels of downsampling (50% to 0.5%).

**Figure S11.**
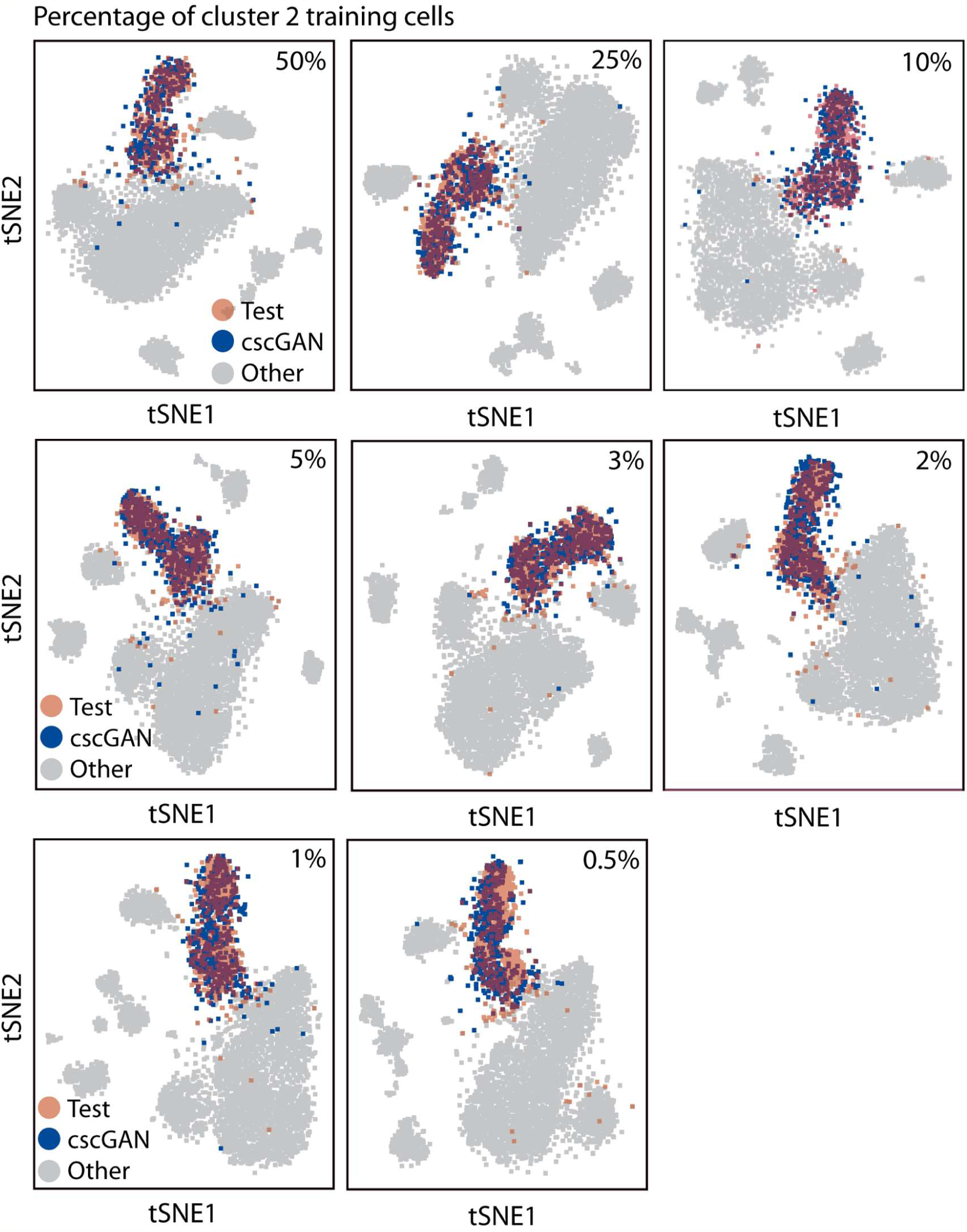
Effect of downsampling on the quality of cscGAN generated cluster 2 cells. Each subfigure is a t-SNE representation of real test (red) and cscGAN generated (blue) cluster 2 cells for different levels of downsampling (50% to 0.5%). Gray cells represent real test data of other clusters.

**Figure S12.**
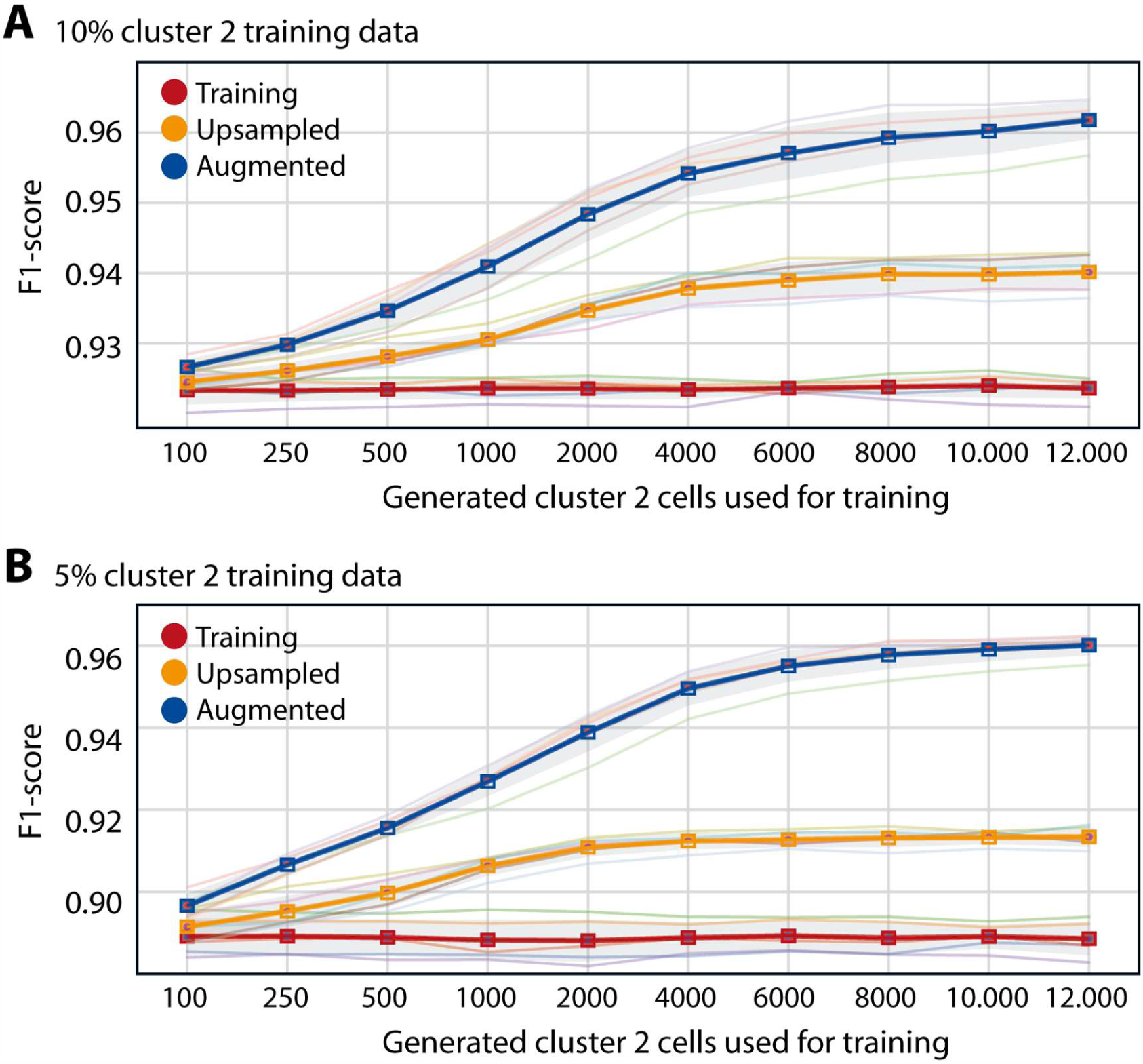
RF classification performance for different numbers of cluster 2 cells. F1-score reached by a RF classifier discriminating cluster 2 from other cells when trained on (A) 10% or (B) 5% downsampled (red), upsampled (yellow), and augmented (blue) cells. The different numbers of upsampled and augmented cells used for training are shown on the x-axis. It is important to note that the number of cluster 2 training cells did not change for the red curve. The results represent the mean for five different data partitions (seeds).

